# Unexpected CRISPR off-target mutation pattern *in vivo* are not typically germline-like

**DOI:** 10.1101/193565

**Authors:** Zhiting Wei, Funan He, Guohui Chuai, Hanhui Ma, Zhixi Su, Qi Liu

## Abstract

Schaefer et al.^1^ (referred to as **Study_1**) recently presented the provocative conclusion that CRISPR-Cas9 nuclease can induce many unexpected off-target mutations across the genome that arise from the sites with poor homology to the gRNA. As Wilson et al.^2^ pointed out, however, the selection of a co-housed mouse as the control is insufficient to attribute the observed mutation differences between the CRISPR-treated mice and control mice. Therefore, the causes of these mutations need to be further investigated. In 2015, Iyer et al.^3^ (referred to as **Study_2**) used Cas9 and a pair of sgRNAs to mutate the Ar gene in ***vivo*** and off-target mutations were investigated by comparison the control mice and the offspring of the modified mice. After analyzing the whole genome sequencing (WGS) of the offspring and the control mice, they claimed that off-target mutations are rare from CRISPR-Cas9 engineering. Notably, their study only focused on indel off-target mutations. We re-analyzed the WGS data of these two studies and detected both single nucleotide variants (SNVs) and indel mutations.

---

Because these two studies draw relatively opposite conclusions on the off-targeting of the CRISPR system on the whole genome, the origins or causes of the mutations need to be cautiously examined. Here we performed a computationally evolutionary investigation (***Figure 1a***) to re-analyze the WGS data of these two studies with a direct comparison of the experiment design and analysis results (***Table 1*** and ***Supplementary Table 1).*** The computational framework designed for the above analysis can accurately infer the likelihood of the origins of these mutations, ***i.e.***, whether these mutations are germline-like or not. This controversial issue arises substantially arguments but is unresolved^1–4^. Here our analysis concluded that the so-called unexpected SNVs pattern (***Supplementary Notes*** and ***Figure 1a***) in **Study_1** are not typically germline-like, while for **Study_2,** the detected SNVs are in fact germline mutations.

**Figure 1.**
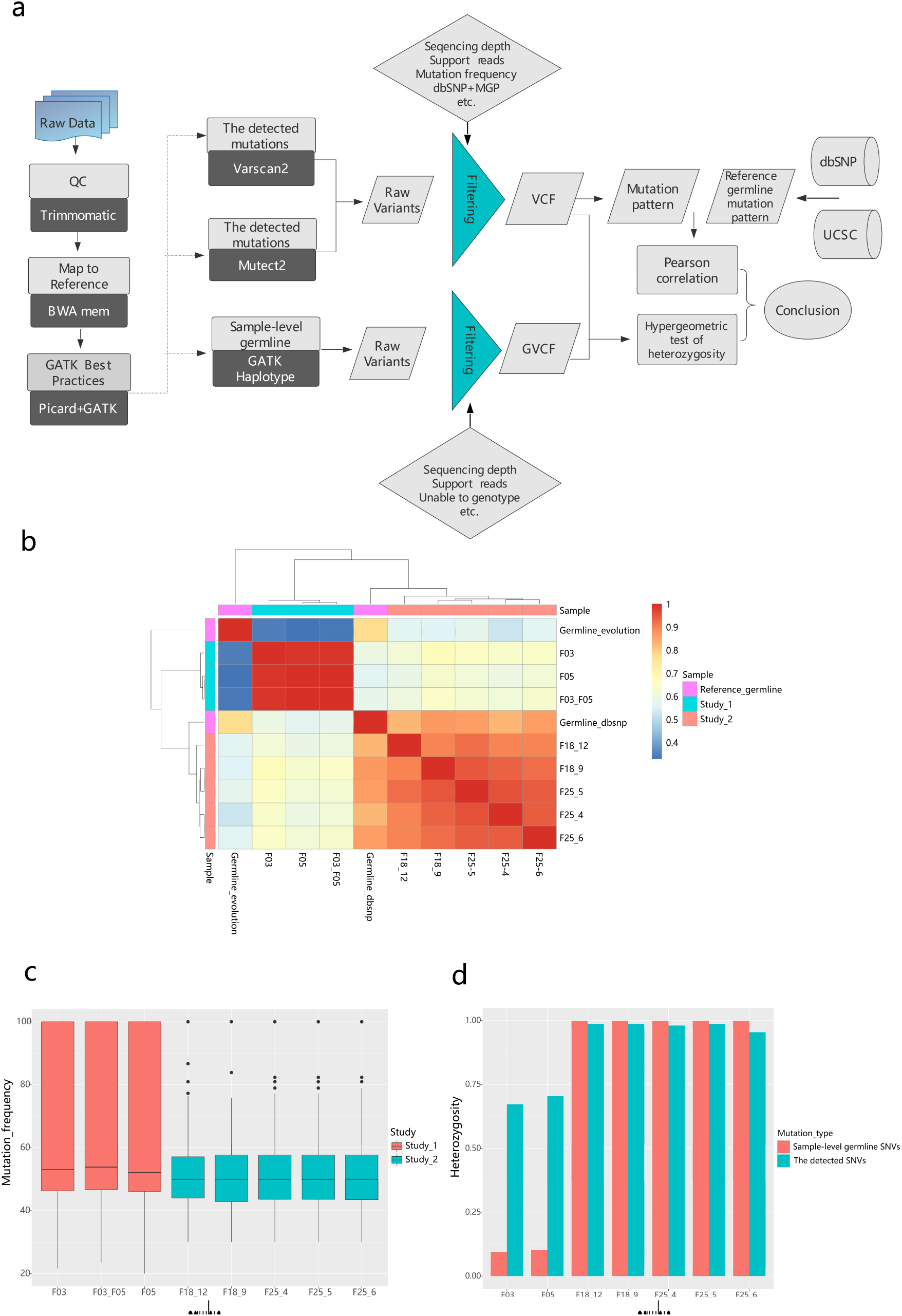
(a) Overview of our computational framework to re-analyze the WGS data of these two studies. (b) Heatmap of the similarities of the mutation pattern between the reference germline SNVs and the detected SNVs. (c) The detected SNVs mutation frequency in these two studies. (F03 and F05 are the samples from Study_1, F03_F05 SNVs is the intersection SNVs of these two samples. F18_9, F18_12, F25_4, F25_5 and F25_6 are the samples from Study_2). (d) Heterozygosity of the detected SNVs and the sample-level germline SNVs.

**Table 1.**
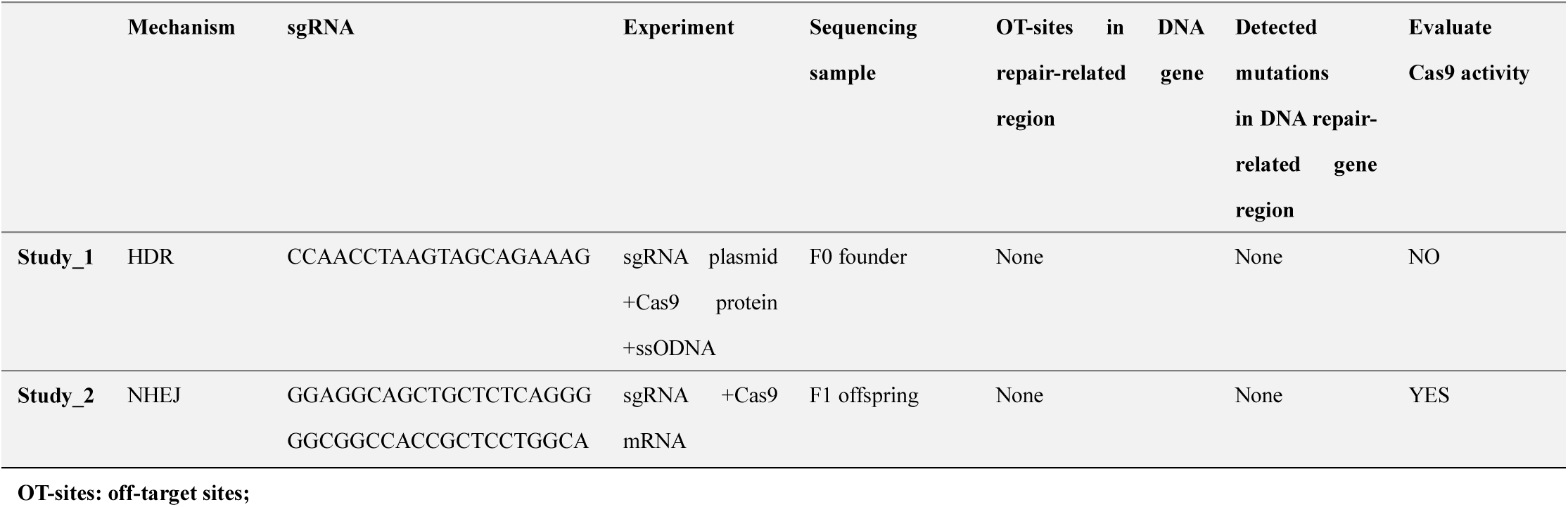
A direct comparison of Study_1 and Study_2.

Thousands of genomic mutations were found from both studies based on our computational framework (***Supplementary Table*** 2). Following our strict criteria to filter SNVs with low mutation frequency and indels overlapping with the UCSC Genome Browser^5,6^ short tandem repeat, we confirmed a low number of so-called unexpected mutations claimed in Study_1**^*1*^(*Supplementary Table*** 2). A great deal of filtered mutations are potentially false positives arising from spontaneous mutations, sequencing and Burrows-Wheeler Aligner(BWA)^7^ alignment errors. This leads us believe that the calling of CRISPR-induced mutations especially indels should be carefully performed to avoid false positives. We integrated two computational methods to infer the putative origins of these detected SNVs, by (1) quantitatively analyzing the similarity of the mutation pattern^8–11^ (***Supplementary Table 3***) between the reference germline SNVs (derived from well-curated public databases^5,9^; ***Figure 1a*** and ***Supplementary Notes***) and the detected SNVs derived from Study_1 and Study_2 followed our re-analysis pipeline. The rationale is that for a particular organism (e.g., mouse), its germline mutation pattern should be evolutionarily conserved; (2) we applied a hypergeometric distribution to test the difference of mutation heterozygosity between the sample-level germline SNVs (detected using ***GATK HaplotypeCaller***; ***Figure 1a*** and ***Supplementary Notes***) and the detected SNVs.

As seen in the heatmap (***Figure 1b***), the detected SNV mutation pattern correctly cluster the samples from different studies. In addition, the mutation pattern of samples from **Study_2** highly correlated with the reference dbSNP germline mutation pattern (R^2^∼0.85, *P-value* <2.2e-16), while the correlation between the samples from **Study_1** and the reference dbSNP germline was weaker (R^2^∼0.55, *P-value*∼1e-10). Although the correlation of mutation pattern in either study with germline at evolution scale are relatively low, we found that the samples from Study_2 correlated stronger with germline mutation pattern than those of samples from **Study_1.**

The heterozygosity of the so-called unexpected SNVs of the samples in **Study_1** was 67.2% and 70.4%, respectively (***Figure 1c***), while that of the sample-level germline SNVs was much lower (***Figure 1c***). We reasoned that if all the detected SNVs were germline, the heterozygosity of such SNVs would be nearly identical to that of the sample-level germline SNVs. Our statistical test, however, indicates a significant difference between the heterozygosity of them (p-value=0; ***Supplementary Notes***), proving that the overall derived mutation pattern in Study_1 is not germline-like. Note that Lareau et al. and Kim et al.^4,6^ recently demonstrated that the two CRISPR-Cas9 treated mice (F03, F05) in **Study_1** are actually more closely related to each other genetically than to the control mouse, proving that these mutations are most likely pre-existing variants. Our analysis result further presented that there are unusual SNVs arose in **Study_1** combined with pre-existing variants, and the overall mutation pattern in vivo are not germline-like. In contrast, in **Study_2**, we cannot reject the null hypothesis that the detected SNVs were sampled from the germline background. We computationally validated that the majority of the detected mutations in **Study_**2 are germline, which is consistent with their original notation that off-target somatic mutations are rare in their Cas9-modified mice^3^.

Because the same CRISPR genome-editing technology led to different conclusions in two studies, we performed a direct comparison of these two studies to evaluate the possible causes for the differences (***Table 1*** and ***Supplementary Table*** 1). First, the experimental protocols differ, where the protocol used in **Study_1** is not a routine and well-accepted CRISPR knockout protocol (sgRNA plasmid + Cas9 protein). The potentiality to induce unusual mutation patterns with such protocol are waiting to be further explored. Secondly, the whole genome sequencing of the founder mouse should be performed before CRISPR knockout. The founder genetic variation should be carefully examined in both studies, since such genetic variation, which may contain in the zygotes, can confound the target sites of certain sgRNAs more than others. This information should be integrated into the study for sgRNA selection to ensure safety^12^.

In summary, we demonstrated that the unexpected CRISPR off-target mutation pattern in **Study_1** are not typically germline-like. Some of unusual and unidentified mutations may arise in **Study_1**, but the real reasons remain to be explored. Based on the available data and a direct comparison of the two studies, we presented two possible reasons and future re-analysis directions that may contribute to such different conclusions. To characterize the authentic CRISPR-mediated mutations, we are required to have appropriate controls to rule out other sources of mutations, which will be needed for benchmarking of targeting safety of CRISPR-based gene therapy.

## Acknowledgments

This work was supported by the National Major Research and Innovation Program of China (Grant No. 2016YFC1303205, SQ2017YFSF090222), National Natural Science Foundation of China (Grant No. 61572361), Shanghai Rising-Star Program (Grant No. 16QA1403900), and Shanghai Natural Science Foundation Program (Grant No. 17ZR1449400).

